# Genomic G quadruplexes regulate mRNA splicing

**DOI:** 10.64898/2025.12.15.693900

**Authors:** Victoria Mauz, Thiago Britto-Borges, Tobias Merkel, Enio Gjerga, Joshua Hartmann, Marteinn Thor Snaebjörnsson, Friederike Schreiter, Christian U. Oeing, Carsten Sticht, Samuel Sossalla, Jürgen Günther Okun, Georg Stoecklin, Matthias W. Hentze, Eileen E. M. Furlong, Almut Schulze, Matthias Dewenter, Christoph Dieterich, Johannes Backs

**Author notes:** Corresponding author: Johannes Backs, MD Heidelberg University, Medical Faculty Heidelberg Im Neuenheimer Feld 669 69120 Heidelberg, Germany. these authors contributed equally (Matthias Dewenter, Christoph Dieterich, Johannes Backs).

## Abstract

Genomic G quadruplexes (G4) are non-canonical DNA structures that regulate gene expression primarily through transcriptional control. Here, we uncover that DNA G quadruplexes are critical determinants of pre-mRNA splicing. G4s on the DNA template strand serve as recruiting elements for the RNA-binding proteins SRSF9 and WBP11 in order to facilitate productive splicing of adjacent pre-mRNAs. This process is controlled by protein arginine methyltransferase 5 (PRMT5) that releases SRSF9 and WBP11 from G4s through arginine methylation. Acyl-CoA dehydrogenase very long-chain specific (*ACADVL*) is a gene highly regulated by this mechanism since targeting G4 stability prevents mis-splicing and restores ACADVL protein levels. In the heart, deletion of *Prmt5* recapitulates defective splicing and results in progressive cardiac failure along with loss of ACADVL in mice. Importantly, we find that *Acadvl* regulation plays a critical role for cardiomyopathy as restoring *Acadvl* expression in *Prmt5* knockout mice prevents cardiac dysfunction. This study establishes an unanticipated fundamental principle by which genomic G quadruplexes act as splicing enhancers of associated pre-mRNAs, and reveals an essential role in cardiac homeostasis.

## Introduction

Co- and post-transcriptional processing of mRNA represents a major source of complexity in eukaryotes by enabling tissue- and cell-specific expression of mRNA and protein isoforms. The precise recognition and removal of introns requires a highly organized and controlled process performed by the spliceosome – a ribonucleoprotein complex of dynamic composition comprising more than 300 proteins^1^, which by themselves are often subject to post-translational modifications. Arginine methylation is known to critically modulate mRNA splicing by tuning the affinity and activity of spliceosomal components and splicing-associated factors^2^. Protein arginine methyltransferase 5 (PRMT5) is an important type II methyltransferase catalyzing mono- (MMA) and symmetric dimethylation of arginine residues (SDMA)^3^. Whereas PRMT5 is required for cellular homeostasis and integrity, altered activity is sufficient to promote malignant transformation, tumor progression and heart failure^4,5^. PRMT5 interacts with and methylates Sm proteins^6^ and splicing factors^7^, thereby promoting spliceosome assembly and activity. As a result, PRMT5 inhibition induces defective pre-mRNA splicing, which is clinically relevant as it provides a therapeutic approach for tumors vulnerable to aberrant splicing^8^.

Despite extensive evidence for its critical regulation of mRNA splicing, the underlying mechanism of PRMT5-dependent splice site selection is not known. Using unbiased structural analyses of retained introns upon PRMT5 inhibition, we uncovered a strong association with the guanine quadruplex motif (G quadruplex, G4) indicative of non-canonical DNA structure formation. G4s are four-stranded, non-canonical structures that can be formed by guanine-rich sequences via selective guanine-guanine hydrogen bonds^9^. Systematic integration of G4-targeted sequencing confirmed a highly significant correlation with G4s on the DNA template strand of the retained intron on a genome-wide scale, suggestive of a globally relevant mechanism. One of the most PRMT5-regulated genes encodes for acyl-CoA dehydrogenase very long-chain specific (ACADVL), an integral component of mitochondrial β-oxidation. Interfering with G4 formation decreased splicing efficiency of *ACADVL*, while selective G4 stabilization enabled efficient splicing and restored ACADVL protein levels. Through an unbiased screen we identified SRSF9 and WBP11 as functional mediators of PRMT5-dependent intron retention (IR), and further demonstrate that the G4-binding properties of SRSF9 and WBP11 are controlled by PRMT5-dependent methylation, suggesting a direct link between arginine methylation and G4-dependent splicing. Since ACADVL deficiency in humans is causative for infantile cardiomyopathy^10^, we used a cardiomyocyte-specific genetic model of PRMT5-deficiency (*Prmt5*-cKO) and confirmed massive retention of G4-associated introns. Restoring ACADVL expression protected *Prmt5*-cKO mice from heart failure, demonstrating pathophysiological relevance of the newly identified mechanism. Collectively, our data suggest that genomic G4s enhance splicing of adjacent pre-mRNAs through PRMT5-dependent binding of SRSF9 and WBP11.

## Results

### Retained introns are enriched at genomic G quadruplexes

In order to gain insights into PRMT5-dependent splicing, we used the substrate-competitive compound EPZ015666 as a specific pharmacological PRMT5 inhibitor (PRMT5i)^11^, and subjected nuclear RNA of HEK293 cells to Oxford Nanopore Technology longread sequencing. In line with previous studies^12,13^, PRMT5i induced major alterations of differential splicing along with intron retention (IR) as the most abundant splicing alteration (**Suppl. Fig. 1A**). Unbiased characterization of retained introns revealed significant differences in length, 5’ splice site strength and GC content as compared to their non-regulated controls (**Suppl. Fig. 1B**). Since we were unable to detect specific enrichment of RNA-binding protein (RBP) motifs, including those of known PRMT5 substrates, we hypothesized that secondary and tertiary structures^14^ might differentiate retained from non-regulated introns. Indeed, unbiased prediction suggested a strong association between retained introns and G quadruplexes (G4) on the DNA template strand, while no enrichment for other non-B helical structures was observed (**Fig. 1A**, **Suppl. Fig. 1C**). G4s represent non-canonical structures composed of guanine tetrads that interact through Hoogsteen base pairing^9^. Despite a G4 consensus motif, computational prediction significantly differs from experimentally validated structures^15^ indicating large variability. We therefore utilized a published resource of experimental G4-sequencing of the human genome^16^. Strikingly, integrative analysis uncovered a strong association between retained introns upon PRMT5i and G4s, specifically on the DNA template strand of retained introns (**Fig. 1b**, **Suppl. Fig. 1D-F**). Of note, the correlation was even stronger when we applied a window of up to 500bp up- and downstream of the G4s, indicating that almost 40 % of retained introns are highly enriched at or adjacent to genomic G4 positions. As control, non-regulated introns with FDR>0.05 or -0.5<log2(FC)<0.5 were used. This result was consistent across all experimental strategies to stabilize G4s via either the small molecule ligand Pyridostatin (PDS), potassium ions or a combination of both. In contrast, the respective non-template DNA strand did not reveal any significant association indicating a strong strand selectivity.

**Figure 1:**
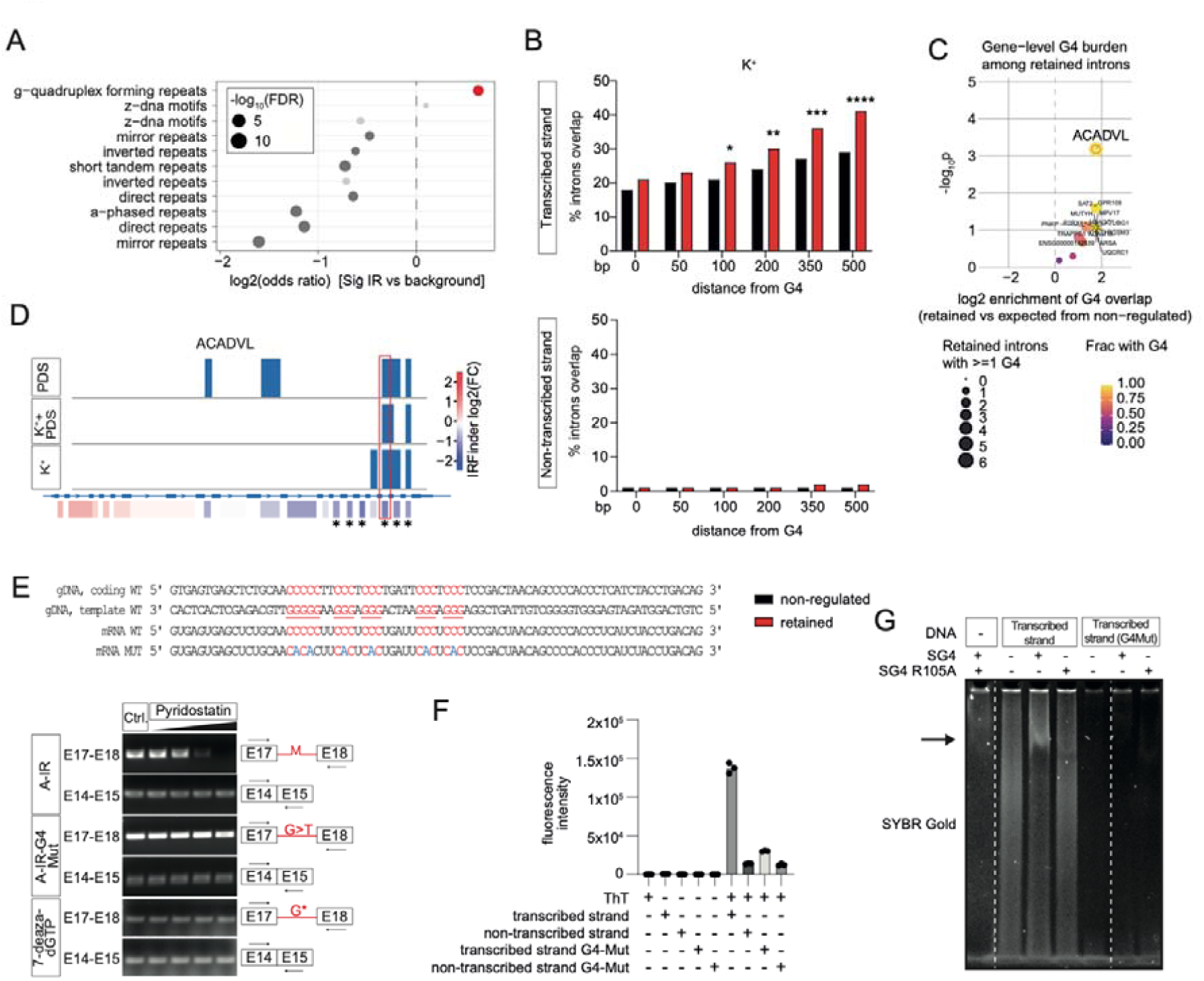
Retained introns are enriched at genomic G quadruplexes. (**A**) Enrichment analysis of non-B DNA motifs among significantly retained introns reveals strong enrichment with G4s on a genome-wide basis using a window of 500bp beyond intron junctions. The point size corresponds to -log10(FDR). Bold and red dots represent FDR<0.05. Extended data sets are depicted in Suppl. Fig. 1c. (**B**) Integrated analysis of G4-seq and ONT-seq in HEK293 nuclei upon PRMT5i. Potassium ions were used as G4 stabilizers. The correlation of retained and non-regulated introns is shown for a window of ±500bp around the G4. Transcribed and non-transcribed DNA strands are separately analyzed. Statistical analysis: * p=0.0458, ** p=0.00621, *** p=0.00012, **** p<0.0001 (Proportion test). Extended data sets are depicted in Suppl. Fig. 1D,E. (**C**) G4 enrichment analysis on gene level highlights ACADVL as prominent target of G4-associated IR using a window of ±500bp. The volcano plot depicts the relative fraction of introns overlapping with at least one G4 as compared to the expected overlap rates in non-regulated introns. Potassium ions were used as G4 stabilizers. Extended data sets are depicted in Suppl. Fig. 1G. (**D**) Tracks of G4-seq across different experimental conditions including Pyridostatin (PDS) and/or potassium ions (K^+^) as G4 stabilizing agents mapping to *ACADVL* (upper panel) and Oxford Nanopore longread nuclear RNA-sequencing upon PRMT5i in HEK293 analyzed by IRFinder (lower panel). IR is depicted by negative log2FC and blue color. Significantly retained introns are marked by asterisks. The genomic region corresponding to intron 17/18 is highlighted. (**E**) Sequences of intron 17/18 on genomic DNA level (gDNA) and transcribed mRNA (upper panel). The predicted G4-forming region is underlined. mRNA G4-Mut includes 6 point mutations shown in blue disrupting G4 stretches. Polymerase stalling assay using either a wildtype or G4-mutated minigene along with increasing dosages of Pyridostatin (lower panel). 7-deaza-dGTP incorporation is used in A-IR and restores amplification. As reference region and loading control a distant amplicon between exon 14 and 15 without predicted G4 formation was used (E14-E15). (**F**) Fluorescence staining of G4s using Thioflavin T (ThT) reveals G4 formation in the transcribed DNA stand of I17/18 which is prevented by G4 mutations (G4-Mut). (**G**) G4-specific nanobody (SG4) binds to the G4 in I17/18, but not to the G4-mutated control as determined by EMSA. SG4 R105A serves as non-specific nanobody control lacking G4 affinity.

Among the genes affected by G4-associated IR, *ACADVL* is one of the most prominent targets (**Fig. 1C, Suppl. Fig. 1G**). *ACADVL* hosted 6 significant IR events, all of which are located at the 3’ terminus in immediate proximity to each other. Consistent with the genome-wide correlation, two retained *ACADVL* introns contain G4s on the DNA strand that serves as template for transcription (**Fig. 1D**). Metagene analysis therefore did not point to significant sequencing bias between experimental conditions, implying biological rather than technical causes for the observed correlation (**Suppl. Fig. 1H**). Interestingly, the predicted G4 in *ACADVL* intron 17/18 was present across a broad range of mammals, suggesting an evolutionary conserved element with potential functional relevance (**Suppl. Fig. 1I**). Given the broad conservation, the murine genomic sequence was applied as reference for further studies. To validate G4 presence in *Acadvl* experimentally, we used Pyridostatin as a small molecule ligand known to stabilize G4s^17^. Pyridostatin dose-dependently induced Taq polymerase stalling during amplification of the putative secondary structure, indicating G4 formation which interferes with enzymatic action (**Fig. 1E**, E17-E18). In contrast, amplification of another region from the same gene without predicted G4 did not show any alteration and serves additionally as loading control (**Fig. 1E**, E14-E15). Disruption of guanine repeats by single point mutations (**Fig. 1E**, mRNA Mut) as well as the incorporation of 7-deaza-dGTP, a guanosine analog which is not able to form Hoogsteen base pairs^18^, reversed Pyridostatin-dependent Taq polymerase stalling (**Fig. 1E**). These results provide evidence that the template strand of intron 17/18 is able to form a G4. As orthogonal approach, Thioflavin T was applied as G4-specific fluorescent sensor which resulted in a specific light up phenomenon when incubated with a synthetic wildtype DNA oligonucleotide of intron 17/18 but not with a G4 mutant or complementary DNA sequences supporting G4 formation (**Fig. 1F**). Third, a G4-specific nanobody^19^ indicated complex formation with the wildtype DNA oligonucleotide, while lacking affinity to a G4-mutated control as determined by EMSA (**Fig. 1G, Suppl. Fig. 1J**). In conclusion, complementary computational prediction and experimental approaches uncovered a strong correlation between the position of genomic G4s and introns retained upon PRMT5i.

### Genomic G quadruplex formation controls *ACADVL* splicing

Genomic G4s are critical regulatory elements particularly in promoters and telomeres, and have been described in the context of transcriptional control as well as genomic integrity^9^. However, a potential relevance for IR has not been described yet. In order to further characterize PRMT5-dependent mechanisms, we first cross-validated the long-read sequencing results via qPCR. PRMT5i induced dose-dependent IR in nuclear RNA, leading to a decrease in productively spliced *ACADVL* (**Fig. 2A**). Interestingly, subcellular fractionation revealed that intron-containing transcripts predominantly accumulate in the nucleus (**Fig. 2A**, **Suppl. Fig. 1K**). In contrast, spliced *ACADVL* mRNA was well detectable in the cytoplasm, suggesting that IR causes nuclear detention of *ACADVL* pre-mRNA, thereby preventing ACADVL protein production. Indeed, western blot analysis confirmed a selective, PRMT5i dose-dependent loss of ACADVL protein (**Fig. 2B**). Blunted palmitate utilization as a read-out of ACADVL activity was measured via Seahorse assays using C16 as primary substrate (**Fig. 2C**) validating that PRMT5i causes a functionally relevant ACADVL deficit.

**Figure 2:**
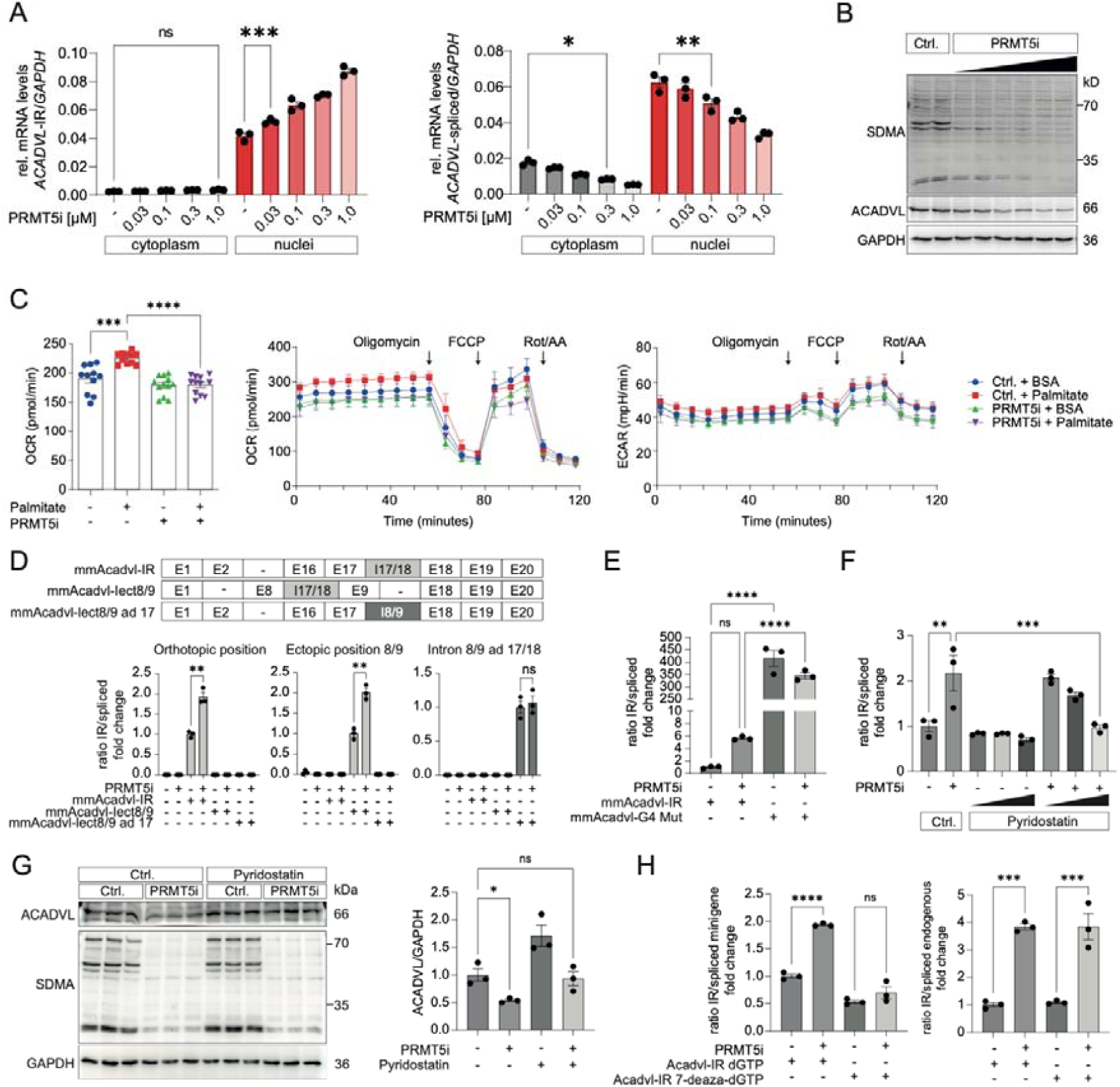
Genomic G quadruplexes control *ACADVL* splicing. (**A**) Quantitative PCR on intron-retained (left), and canonically spliced (right) transcripts of *ACADVL* in nuclear and cytoplasmatic extracts of HEK293 cells upon PRMT5 inhibition (PRMT5i). Values were normalized to *Gapdh*. Statistical analysis: values are presented as Mean±SEM (left) *** p=0.0005 (Ordinary one-way ANOVA and Tukey‘s multiple comparisons test) (right) * p=0.0203, ** p=0.0023 (Ordinary one-way ANOVA and Tukey‘s multiple comparisons test). (**B**) Immunoblot analysis of HEK293 cells treated with PRMT5i. PRMT5i was examined by reduced levels of overall symmetrically dimethylated arginine (SDMA). GAPDH served as loading control. (**C**) Seahorse analysis of HEK293 cells treated with PRMT5i and exposed to palmitate or BSA control. (C) indicates basal respiration values as determined by Agilent’s Wave software. Statistical analysis: values are presented as Mean±SEM, *** p=0.0001, **** p<0.0001 (Ordinary one-way ANOVA and Tukey‘s multiple comparisons test). (**D**) Schematic illustration of the minigene constructs consisting of all 20 exons (E) of the murine reference transcript extended by intron 17/18 (I17/18) either at its natural position (mmAcadvl-IR) or ectopically expressed between exons 8 and 9 (mmAcadvl-Iect8/9). The third construct comprises intron 8/9 expressed between exons 17/18 (mmAcadvl-I8/9 ad 17/18). Lower panel: Quantitative PCR on intron-retained and canonically spliced transcripts of the minigene constructs depicted in upper panel in nuclear extracts of HEK293 cells upon PRMT5i. Values are shown as ratio. Statistical analysis: values are presented as Mean±SEM ** p=0.0015 (Orthotopic position) ** p=0.0016 (Ectopic position 8/9) (Unpaired t test). (**E**) Quantitative PCR on intron-retained and canonically spliced transcripts of minigene constructs in nuclear extracts of HEK293 cells upon PRMT5i. mmAcadvl-G4Mut thereby corresponds to the mutated sequence displayed in Fig. 1d. Values are shown as ratio. Statistical analysis: values are presented as Mean±SEM **** p<0.0001 (Ordinary one-way ANOVA and Tukey‘s multiple comparisons test). (**F**) Quantitative PCR on intron-retained and canonically spliced transcripts of the wildtype minigene in nuclear extracts of HEK293 cells upon PRMT5i and increasing dosages of Pyridostatin. Values are shown as ratio. Statistical analysis: values are presented as Mean±SEM ** p=0.0011 *** p=0.0008 (Ordinary one-way ANOVA and Tukey‘s multiple comparisons test). (**G**) Immunoblot analysis and quantification of HEK293 cells treated with PRMT5i and/or Pyridostatin. Sufficient PRMT5i was validated by reduced levels of overall symmetrically dimethylated arginine (SDMA). GAPDH served as loading control. Statistical analysis: values are presented as Mean±SEM, * p=0.0415 (Ordinary one-way ANOVA and Tukey‘s multiple comparisons test). (**H**) Quantitative PCR on intron-retained and canonically spliced transcripts of the wildtype minigene consisting of either canonical dGTP or 7-deaza-dGTP. Values are shown as ratio. Endogenous human ACADVL served as control for equal PRMT5i (right panel). Statistical analysis: values are presented as Mean±SEM **** p<0.0001 (minigene) *** p=0.0002 (dGTP, endogenous) *** p=0.0002 (7-deaza-dGTP, endogenous) (Ordinary one-way ANOVA and Tukey‘s multiple comparisons test).

Next, we established an *Acadvl* minigene construct in order to address the mechanism of PRMT5i-induced IR (**Fig. 2D**, scheme). Taking advantage of the murine sequence allows to distinguish exogenous *Acadvl* from endogenous human *ACADVL* when expressed in HEK293 cells, whereby functional effects can be assigned unambiguously to the minigene construct. Recapitulating endogenous *ACADVL*, the minigene exhibited increased IR upon PRMT5i as determined by the ratio of intron-containing to spliced transcripts (**Fig. 2D**, orthotopic position). To delineate potential exonic from intronic regulatory elements, we modified the minigene in such way that intron 17/18 is expressed ectopically between exons 8 and 9 whose natural adjacent intron is not subject of PRMT5-dependent regulation as determined by long-read sequencing. In addition, intron 8/9 (88nt) has a similar length to intron 17/18 (78nt). Interestingly, intron 17/18 was retained to similar extent in this ectopic context (**Fig. 2D**, ectopic position 8/9). Conversely, intron 8/9 expressed between exons 17 and 18 did not show any PRMT5-dependent regulation (**Fig. 2D**, Intron 8/9 ad 17). Taken together, these results indicate that IR in *Acadvl* is determined by the intronic sequence itself and not by regulatory elements in the surrounding exons further supporting the hypothesis of a G4-mediated mechanism.

To test effects of G4 perturbations, we introduced point mutations disrupting G stretches in order to prevent G4 formation. Transfected into cells, the G4 mutant exhibited abolished splicing of intron 17/18 as reflected by a >100fold increase of the IR/spliced ratio and the absence of correctly spliced transcripts (**Fig. 2E**). Guided by these data, we next set out to address if the G4 might be functionally involved in PRMT5-dependent *Acadvl* splicing. Combining PRMT5i with Pyridostatin, a well-established G4 stabilizer, we found that IR could be dose-dependently reversed by Pyridostatin (**Fig. 2F**). The *Acadvl* construct allows to relate the mechanistic effect of Pyridostatin specifically to intron 17/18 since it contains the only predicted G4 structure in this minigene. While PRMT5i induced loss of ACADVL protein expression as shown above, ACADVL protein levels were preserved when PRMT5i was combined with Pyridostatin treatment (**Fig. 2G**), indicating that genomic G4 is critical for PRMT5-dependent *ACADVL* splicing and protein production. Similarly, a minigene harboring 7-deaza guanine instead of guanine was not regulated by PRMT5 (**Fig. 2H**) validating that genomic G4 is critical for PRMT5-dependent splicing regulation.

### SRSF9 and WBP11 are critical mediators of G4-dependent splicing

We then turned our focus towards the methyltransferase activity, the second component required for PRMT5-dependent splicing. Overall, PRMT5 is known to interact with and methylate dozens of RNA-binding proteins (RBPs), which have commonly been evaluated as splicing regulators^7^. Given the importance of genomic G4s in determining splicing outcome, we reasoned that methylation substrates relevant for pre-mRNA splicing might be associated with genomic DNA beyond their role as RBPs.

To identify such proteins through an unbiased approach, we conducted a functional screen based on shRNA-mediated knockdown of 87 proteins known as arginine methylation targets with nucleic acid binding capacity (**Suppl. Table 1**). Thereby we found that knockdown of SRSF9, WBP11, ILF3, MECP2 and CHTOP consistently induced *Acadvl*-IR, thereby phenocopying PRMT5i (**Fig. 3A, B**). Among these candidates, knockdown of SRSF9 and WBP11 also led to a decrease in ACADVL protein abundance (**Fig. 3C**), indicating that mis-splicing might be causative for reduced ACADVL expression as observed with PRMT5i. At the same time, SRSF9 and WBP11 were found to be methylated by PRMT5, whereas no differential methylation was detected at other factors such as ILF3 (**Fig. 3D-G**). Of note, PRMT5i led to a >90% reduction of SDMA-modified SRSF9, suggesting that PRMT5 is the main methyltransferase for SRSF9.

**Figure 3:**
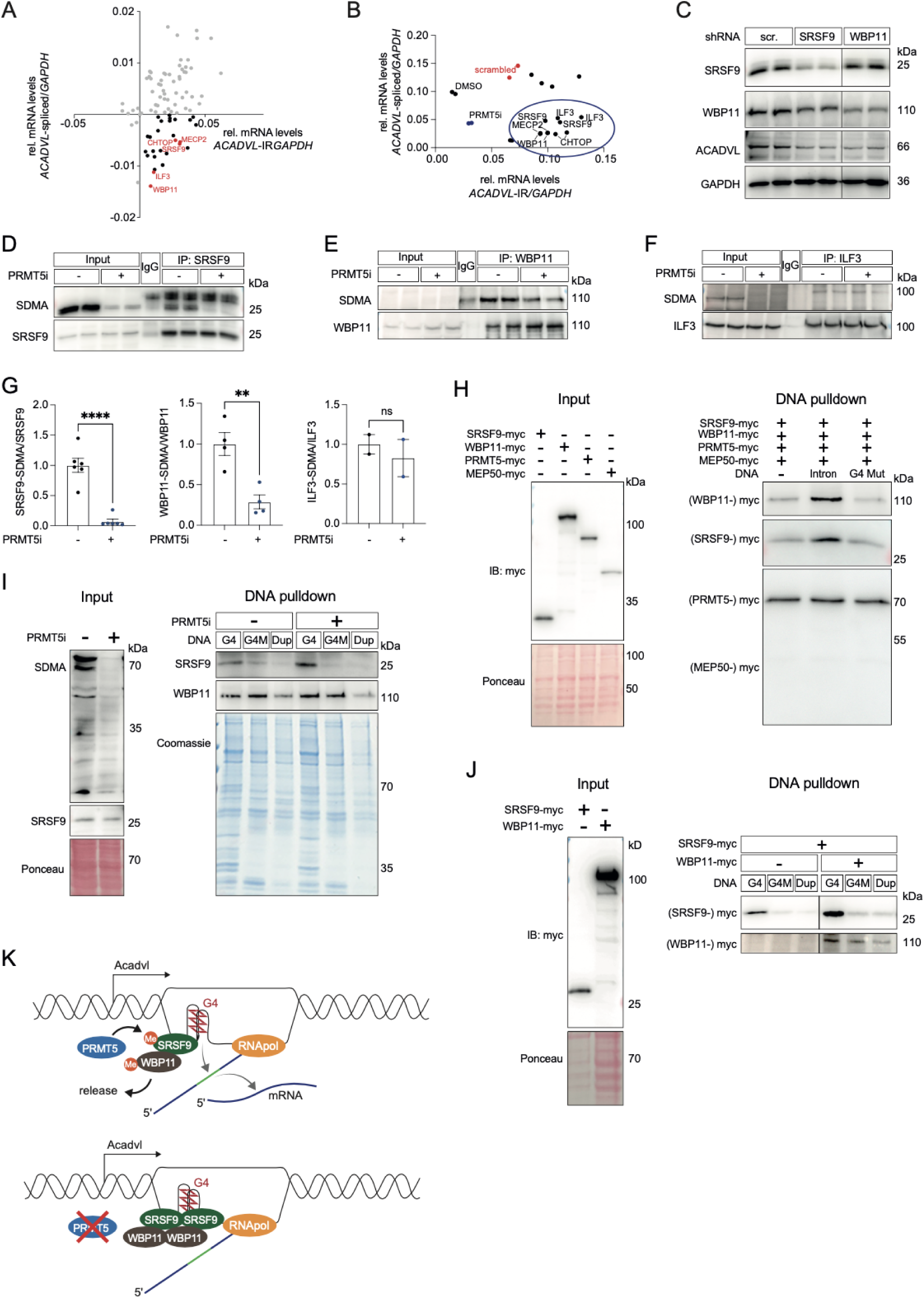
SRSF9 and WBP11 are critical G4-binding proteins mediating dysfunctional splicing. (**A**) Screening approach using quantitative splicing PCR on *ACADVL* in nuclear extracts of HEK293 cells upon shRNA-knockdown of RNA-binding proteins as normalized to scrambled control (cross of x and y axis). Targets inducing increased IR and less productively spliced *ACADVL* are highlighted. Values were normalized to *GAPDH*. (**B**) Quantitative splicing PCR on *ACADVL* in nuclear extracts of HEK293 cells upon knockdown of RNA-binding proteins as compared to scrambled control. Phenocopying PRMT5i, knockdown of SRSF9, WBP11, ILF3, MECP2 and CHTOP induces increased IR and less productively spliced *ACADVL*. Values were normalized to *GAPDH*. (**C**) Immunoblot analysis of HEK293 cells upon knockdown of SRSF9 and WBP11. Interestingly, reduced expression of SRSF9 and WBP11 also repressed ACADVL protein levels. GAPDH served as loading control. Corresponding bands are on the same membrane. (**D-G**) Immunoblot analysis of immunoprecipitation (IP) of endogenous SRSF9 (**D**), WBP11 (**E**) and ILF3 (**F**). SDMA antibody detects reduced methylation of SRSF9 and WBP11 upon PRMT5i as quantified in (**G**) based on independent experiments. (**H**) G4-DNA pulldown upon incubation with myc-tagged candidate proteins expressed in HEK293 cells. A G4-mutated sequence as well as a condition lacking DNA served as negative controls. WBP11 and SRSF9 showed increased affinity to G4 in contrast to PRMT5 and its core partner MEP50. (**I**) G4-DNA pulldown using HEK293 lysates upon PRMT5i. Endogenous SRSF9 exhibits increased binding to G4 upon PRMT5i. A G4 DNA mutant (identical to (H)) and the corresponding G4 DNA double strand (Dup) were used to control for structure-specific binding. Coomassie staining indicates that PRMT5i-induced increased binding to G4 is specific for SRSF9, and does not affect overall protein binding. (**J**) G4-DNA pulldown upon incubation with myc-tagged SRSF9 and WBP11 expressed in HEK293 cells. Controls correspond to (I). SRSF9 exhibits increased binding to G4 DNA in the presence of WBP11. Corresponding bands are on the same membrane. (**K**) Current working model illustrating that PRMT5 is required for productive pre-mRNA splicing of *ACADVL* via controlling differential binding of SRSF9 and WBP11 to genomic G4s.

Given their functional impact and methylation status, we speculated that SRSF9 and WBP11 might also be associated with G4 structures. In order to test this hypothesis, biotinylated G4 DNA and a mutant control were incubated with cellular lysate upon overexpression of either SRSF9, WBP11, PRMT5, or MEP50, a core component of the PRMT5 complex. Indeed, both, SRSF9 and WBP11 were identified as G4 DNA-binding proteins while lacking affinity to G4 mutant DNA (**Fig. 3H**). To address the consequences of PRMT5-dependent methylation, endogenous proteins were used upon PRMT5i or control. PRMT5i enhanced binding of SRSF9 to DNA G4, but not to a G4 mutant nor to the G4-corresponding DNA double strand (Duplex, Dup) that allows to relate SRSF9 affinity to the G4 structure and not to the linear guanin-rich sequence (**Fig. 3I**). In contrast, endogenous WBP11 showed minor PRMT5-dependent effects along with less structure specificity. Comparative binding assays of SRSF9 and WBP11, however, revealed increased G4 binding of SRSF9 when co-incubated with WBP11 suggesting that WBP11 tunes G4 DNA affinity of SRSF9 (**Fig. 3J**).

These data provide a fundamentally new concept of genomic G4s as critical determinants of pre-mRNA splicing. We found that PRMT5-dependent splicing requires G4 formation on the DNA template strand, through a mechanism mediated by SRSF9 and WBP11 as G4 DNA-binding proteins and direct methylation substrates of PRMT5. Methylation of SRSF9 and WBP11 controls their release from the G4 while PRMT5i induces genomic accumulation along with deficient mRNA splicing (**Fig. 3K**).

### G4-mediated ACADVL regulation is critical for cardiac function

ACADVL loss-of-function in humans causes a severe form of infantile cardiomyopathy as clinical manifestation of a rare genetic syndrome (OMIM #201475)^10^. Therefore, we selected the heart as organ where the PRMT5-ACADVL axis might be functionally relevant *in vivo*, and generated a mouse model harboring a cardiomyocyte-specific and inducible knockout (cKO) of *Prmt5* (**Suppl. Fig. 2A**). Cre-positive *Prmt5* wildtype mice in the same genetic background served as controls (*Ctrl.*). Via western blot analysis, a cardiac-specific loss of PRMT5 was validated after cKO induction (**Suppl. Fig. 2B**). PRMT5 therefore seems to be predominantly expressed in cardiomyocytes relative to non-myocytes, as reflected by strongly reduced protein levels in left ventricular tissue. First, the spontaneous phenotype was investigated. *Prmt5*-cKO mice developed progressive cardiac dysfunction within 6 months after cKO induction (data will be reported somewhere else) along with loss of ACADVL (**Fig. 4A**), which recapitulates reduced ACADVL expression upon PRMT5i in HEK293 cells (**Fig. 2B**). Targeted and untargeted lipidomics in cardiac tissue revealed an accumulation of long-chain acylcarnitines (particularly C16 and C18) as primary ACADVL substrates, along with significantly altered triglyceride composition (**Fig. 4B**, **Suppl. Fig. 2C, D**).

**Figure 4:**
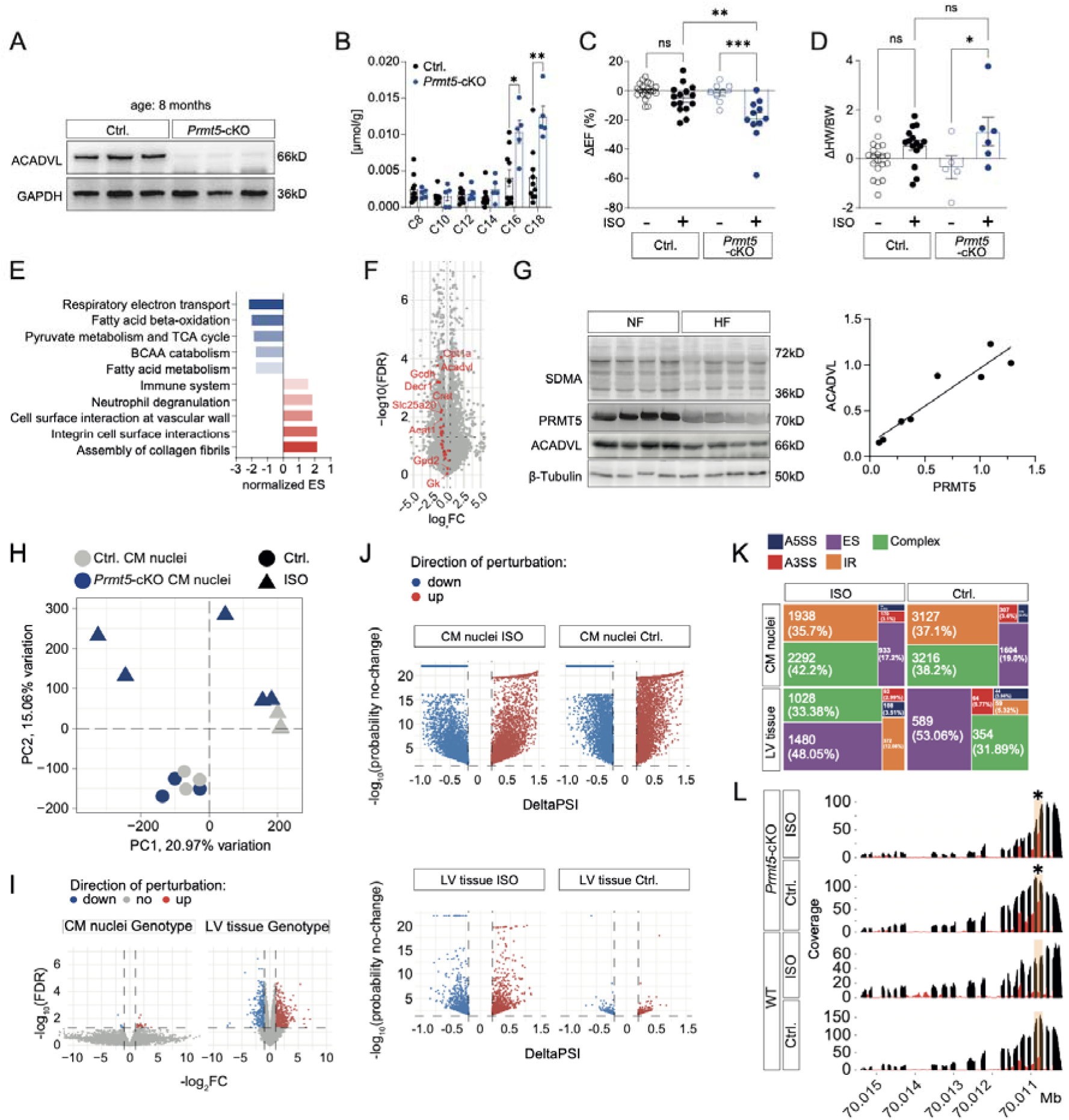
Loss of PRMT5 in cardiomyocytes induces heart failure. (**A**) Immunoblot analysis of cardiac tissue of aged *Prmt5*-cKO mice (6 months after cKO induction, equal to a final age of 8 months) shows loss of ACADVL as compared to controls. GAPDH served as loading control. (**B**) Acyl-carnitine profiling of cardiac tissue reveals accumulation of ACADVL substrates C16 and C18 in aged mice. (**C**) Echocardiographic analysis of *Prmt5*-cKO and control mice after Isoprenaline (ISO) treatment for 10 days. Statistical analysis: ejection fraction (EF) values are presented as Mean±SEM along with Ctrl. without ISO as reference, ** p=0.0023, *** p=0.0008 (Ordinary one-way ANOVA and Tukey‘s multiple comparisons test). (**D**) Analysis of heart weight (HW) of *Prmt5*-cKO and control mice after 10 days of ISO treatment. Values were normalized to body weight (BW). Statistical analysis: values are presented as Mean±SEM, * p=0.0432 (Ordinary one-way ANOVA and Tukey‘s multiple comparisons test). (**E**) Pathway analysis of RNA sequencing of cardiac tissue of *Prmt5*-cKO and control mice after 3 days of ISO stimulation. Pathways were ranked according to their normalized enrichment score (ES). (**F**) Volcano plot of differentially expressed genes highlighting genes compromising the fatty acid metabolism pathway. (**G**) Immunoblot analysis of cardiac tissue of heart failure patients (HF) and healthy controls (NF). β-tubulin served as loading control. Right panel: Correlation between PRMT5 and ACADVL based on immunoblotting shown in left panel and simple linear regression. (**H**) Principal component analysis on RNAseq of cardiomyocyte-specific (CM) nuclei of PRMT5-deficient and control (Ctrl.) mice after 3 days of ISO or NaCl as control. (**I**) Volcano plot depicting differential gene expression analysis of PRMT5-deficient CM nuclei and LV tissue. (**J**) Effect size distribution of alternative splicing events in PRMT5-deficient CM nuclei and LV tissue. (**K**) Size-proportional representation of absolute numbers of alternative splicing types in CM nuclei and LV tissue. We distinguish intron retention (IR), exon skipping (ES), alternative 3‘ splice site (A3SS), alternative 5‘ splice site (A5SS) as well as complex events. (**L**) Alignment coverage plot of the *Acadvl* locus normalized to 500 reads per sample identifying IR in CM nuclei upon *Prmt5*-cKO. Exon and intronic coverage are depicted in black and red, respectively. The highlighted intron maps to chr11:70,010,771-70,010,848 (GRCm38). IR with deltaPSI=0,24 and 0,47 and no-change probability of <0.01 for the ISO and Ctrl. condition, respectively. n≥3.

To accelerate pathological cardiac remodeling, we subjected the mice to daily intraperitoneal injections of isoprenaline (ISO), a β-adrenergic agonist that stimulates cardiac lipolysis and energy production and that is frequently used to induce pathological cardiac remodeling^20–22^. Treatment was started three weeks after cKO induction when cardiac function was still preserved. Interestingly, ISO led to a significantly reduced cardiac ejection fraction within 10 days of treatment in *Prmt5*-cKO mice along with cardiac hypertrophy^23,24^, indicating that loss of PRMT5 decreases the compensatory capacity of the heart in adapting to β-adrenergic stimulation (**Fig. 4C, D**, **Suppl. Table 2**). RNA-sequencing (RNAseq) of heart tissue was conducted three days after ISO stimulation, before cardiac dysfunction was overt. Gene expression analysis revealed 1,926 significant differentially expressed genes, with 57% being downregulated. Gene set enrichment analysis identified pathways associated with mitochondrial energy metabolism to be the most downregulated ones in *Prmt5*-cKO with ISO stimulation, with strong representation of genes in respiratory electron transport, pyruvate metabolism and tricarboxylic acid cycle (TCA) as well as fatty acid oxidation (FAO) (**Fig. 4E, F**). In contrast, inflammatory processes, especially immune cell signaling, and fibrotic remodeling, were upregulated in *Prmt5*-cKO mice. Remarkably, one of the most prominently and consistently regulated genes in the data set encodes for ACADVL, validating the functional relevance of PRMT5 for mitochondrial β-oxidation *in vivo* (**Fig. 4F**).

To investigate whether PRMT5 and ACADVL relate to each other also in patients, we conducted western blot analyses from biopsies of human left-ventricular (LV) myocardium from failing hearts, in comparison to non-failing heart tissue from non-transplanted donor organs. Notably, both PRMT5 and ACADVL levels were strongly reduced in diseased hearts, together with an overall decrease of SDMA as an indicator of reduced type II PRMT activity (**Fig. 4G**). These results point to the relevance of the G4- and PRMT5-dependent splicing mechanism for human cardiac physiology.

To gain deeper mechanistic insights, we conducted RNAseq of cardiomyocyte-specific nuclei (nucCM) in order to deduce 1) cardiomyocyte-intrinsic processes and 2) primary nuclear alterations in comparison to the RNAseq in LV tissue described above (**Fig. 4H**, **Suppl. Fig. 2E, F, Suppl. Table 3**). As for the LV tissue data set, we used a time point after 3 days of ISO treatment, before manifestation of heart failure. While we found 856 differentially expressed genes in the LV tissue *Prmt5*-cKO ISO data set, RNA-seq in nuclei identified only 30 significantly differentially expressed genes (**Fig. 4I**). In contrast, differential RNA splicing analyses revealed a high number of genes with significant differences in alternative splicing (7750, 11320, 2172 and 389 for CM nuclei ISO, CM nuclei control, LV tissue ISO and LV tissue control, respectively, **Fig. 4J**). Alternative splicing events fall into several classes, whereby pronounced differences were observed between CM nuclei and LV tissue. Consistent with our initial results in HEK293 cells, IR events were the predominating type in nuclear RNA libraries (IR: 1938 and 3127, for ISO and control, respectively), while exon skipping (ES) events were the primary type in LV tissue (ES: 1480 and 589 for ISO and control, respectively, **Fig. 4K**). Strikingly, *Acadvl* intron 17/18 harboring the G4 structure was significantly retained by 24% and 47% in *Prmt5*-cKO animals for the ISO and control condition, respectively (**Fig. 4L**, **Suppl. Fig. 2G**), thereby recapitulating the initial cell culture model. An additional IR event in the mouse data set involves *Acadvl* intron 3/4.

### Restoring ACADVL expression prevents heart failure

We identified PRMT5 as a master regulator of cardiac metabolism, in particular mitochondrial β-oxidation, by governing *Acadvl* expression. The identified molecular alterations precede cardiac dysfunction, pointing to a causative role in metabolic remodeling that leads to heart failure^25^. Although we chose to study the role of PRMT5 in the heart due to the described role of ACADVL in rare infantile cardiomyopathy, our data suggest that ACADVL might also play a key role in common forms of heart failure, and thus open the avenue towards novel therapeutic strategies. Experimentally, *Acadvl*-KO mice develop progressive heart failure with reduced ejection fraction (HFrEF)^26,27^. In order to assess the importance of ACADVL dysregulation, AAV9-mediated ACADVL gene transfer was used to compensate the loss of ACADVL in *Prmt5*-cKO cardiomyocytes. Given the mechanistic relevance of genomic G quadruplex formation for PRMT5 regulation, we used a murine *Acadvl* cDNA lacking the intronic G4, and expressed it under the control of the troponin T (Tnnt-) promoter (AAV-Acadvl). A corresponding expression construct encoding renilla luciferase (AAV-Luc) served as a control. Our aim was to restore endogenous ACADVL protein levels as in control animals, and avoid ACADVL overexpression. The appropriate AAV-Acadvl titer was determined in a pilot experiment in *Prmt5*-cKO animals (**Suppl. Fig. 3A-C**). AAV-Acadvl was then administered in a preventive gene therapy approach. Following *Prmt5*-cKO induction, animals were subjected to ISO as described above. *Prmt5*-cKO as well as AAV-mediated expression of ACADVL were confirmed by western blot analysis (**Fig. 5A, B**). Indeed, *Acadvl* levels were equal in control and *Prmt5*-cKO animals, highlighting the regulatory importance of genomic G4. Whereas animals that had received AAV-Luc developed ISO-induced cardiac dysfunction consistent with the previous studies, restoring ACADVL expression was sufficient to prevent HFrEF in *Prmt5*-cKO animals (**Fig. 5C**).

**Figure 5:**
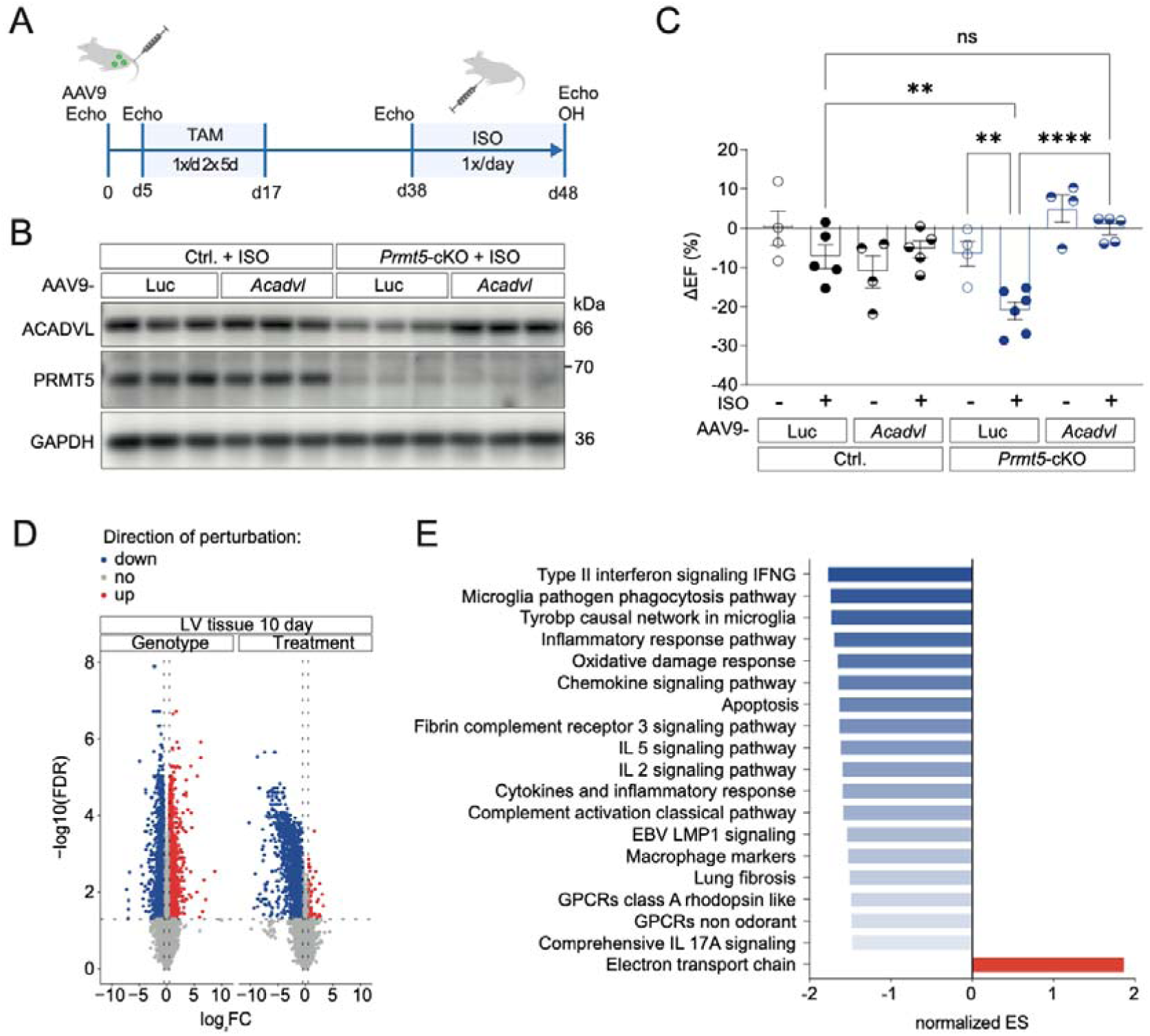
Canonical ACADVL protects from cardiac dysfunction. (**A**) Experimental course of the preventive *Acadvl*-AAV9 gene therapy approach including AAV9 injection, KO induction via Tamoxifen and a 10-days ISO treatment. As control a luciferase-expressing contruct (AAV9-Luc) was applied. (**B**) Immunoblot analysis of ACADVL and PRMT5 to validate viral expression and KO-induction of *Prmt5*-cKO and control mice (Ctrl.) after 10 days ISO treatment. (**C**) Echocardiographic analysis of *Prmt5*-cKO and control mice after AAV9 injection and ISO treatment for 10 days. Statistical analysis: ejection fraction (EF) values are presented as Mean±SEM along with *Ctrl.* Luc without ISO as reference, ** p=0.0043 (*Ctrl.* Luc ISO vs. *Prmt5*-cKO Luc ISO), ** p=0.0051 (*Prmt5*-cKO Luc without ISO vs. *Prmt5*-cKO Luc ISO), p<0.0001 (Ordinary one-way ANOVA and Tukey‘s multiple comparisons test). (**D**) Volcano plot depicting differential gene expression depending on *Prmt5*-cKO (genotype) and ISO (treatment). (**E**) Pathway analysis of RNA sequencing of cardiac tissue of *Prmt5*-cKO treated with Acadvl-AAV compared to Luc-AAV after 10 days of ISO stimulation. Pathways were ranked according to their normalized enrichment score (ES) along with padj <0.05.

To further characterize the cardioprotective function of ACADVL, RNAseq of heart tissue was conducted after 10 days of ISO stimulation comparing the AAV-Acadvl treatment and *Prmt5*-cKO genotype to their respective controls (**Suppl. Fig. 3D**). Whereas *Prmt5*-cKO-dependent alterations included a similar distribution of differentially expressed genes (2164 significantly expressed genes, with 53% being downregulated), AAV-Acadvl transduction induced a strong shift towards gene repression (1366 significantly expressed genes, with 94% being downregulated (**Fig. 5D**). Gene set enrichment analysis identified cytokines and inflammatory signaling as the pathways most affected by AAV-Acadvl administration (**Fig. 5E**, **Suppl. Fig. 3E-G**), indicating ACADVL as a potential hub controlling metabolic and inflammatory pathways.

Taken together, our results provide evidence that loss of ACADVL expression is the underlying cause for *Prmt5*-cKO-induced HFrEF. Given the broad range of metabolic alterations induced by a *Prmt5*-cKO including branched-chain amino acid (BCAA) utilization, Tricarboxylic acid cycle, oxidative phosphorylation and fatty acid oxidation, the identification of ACADVL loss as the single critical mediator of cardiac dysfunction is remarkable, and relates the cardioprotective function of ACADVL to an important role in suppressing inflammation.

## Discussion

### G4 DNA-mediated IR as a novel and general regulatory concept

Genomic G4s belong to non-B helical secondary DNA structures that have so far been characterized primarily in promoters and telomeres^28^. Despite evidence for their biological relevance for transcriptional control and genome stability, their impact on mRNA processing is not known. Our data provide first evidence that G quadruplexes in the genome act as splicing regulators by recruiting SRSF9 and WBP11 to adjacent pre-mRNAs, uncovering a specific DNA feature that controls pre-mRNA processing and localization. This mechanism is critically determined by PRMT5-dependent arginine methylation that enables SRSF9 and WBP11 to be released from G4s. Notably, G4s can also form in RNA, especially within the 5’ UTR of mRNAs where they have been linked to translational control, mRNA localization and splicing^9^. In contrast, our genome-wide approach revealed that only G4s in the template DNA strand (antisense to the coding strand) are linked to PRMT5-dependent splicing, representing a so far unknown mechanism of DNA-dependent splicing activation.

Interestingly, retained introns were found to have an increased GC content^29^ suggesting that genomic G4s may be the underlying reason for this nucleotide bias. We further noticed that PRMT5i-dependent IR events are highly enriched at or adjacent to genomic G4 positions (window ±500bp) as illustrated by *ACADVL*. This raises the possibility that G4s may act by recruiting proteins such as SRSF9/WBP11, interfering with the splicing process of several adjacent introns and physically trapping unspliced transcripts at their genomic origin. Interestingly, a recent study relates PRMT5i-dependently detained transcripts to Sm protein accumulation on chromatin^30^. Indeed, PRMT5i increased binding of SRSF9 to DNA G4s indicative of SDMA that releases protein-DNA G4 interactions. Notably, this model is in good agreement with the observation that arginine (which is positively charged under physiological conditions) not only interacts with the negatively charged phosphate of nucleic acid backbones, but has been shown to bind to guanine through hydrogen bonds which conveys sequence and structure specificity, respectively^31^. Intriguingly, this interaction involves the so called Hoogsteen edge of guanine which is essentially required for Hoogsteen base pairing and, hence, G4 formation^32^. Differential binding of SRSF9 and WBP11 upon PRMT5-dependent SDMA therefore likely interferes with G4 structure formation suggesting a mutual interplay between PRMT5-controlled protein affinity, G4 structure formation and pre-mRNA splicing dynamics.

Furthermore, it will be interesting to examine whether G4 formation on the DNA template strand affects RNA polymerase II, whose elongation speed is tightly connected to splicing efficiency^33^. Preliminary insights from our long read-sequencing experiments do not support the hypothesis that RNA polymerase II would stall at G4 loci. Moreover, G4s seem to be located predominantly in the center of IR hotspots as illustrated by *ACADVL*, while kinetic interference would rather lead to distinct effects up- and downstream of G4s.

### PRMT5 is a master regulator of mitochondrial β-oxidation

Protein arginine methylation is increasingly recognized as a post-translational modification that regulates diverse biological processes including cellular differentiation and proliferation^34^. In line with this notion, PRMT5 was characterized as an oncogene whose gain-of-function correlates with mortality^35,36^ in different cancer entities^37–39^. Therefore, PRMT5 is a promising target in cancer therapy, which has led to the development of enzymatic inhibitors^11^. Under physiological conditions, PRMT5 is essential for the development and homeostasis of various tissues including germ cells, neurons and cardiomyocytes^4,40,41^, and has been implicated in heart failure^5,42^. Our data strongly suggest that cardiac PRMT5 is a key driver of genes related to mitochondrial energy metabolism, especially FAO in cardiomyocytes. Among the genes regulated by PRMT5, ACADVL is essential for maintaining the energy supply in cardiomyocytes, particularly during adaptation to β-adrenergic stress. Recently, PRMT5 was attributed a cardioprotective role in fibroblasts^43^, indicative of a broader role in the cellular cross-talk in complex organs and tissues. While PRMT5 is likely to have cell type-specific functions, our work suggests that PRMT5 regulates conserved downstream effectors (such as ACADVL) in both cardiomyocytes and proliferating cells (HEK293). We propose that differential interaction with G quadruplexes is an immediate downstream effect of PRMT5-dependent methylation, which might not only regulate pre-mRNA splicing but also gene expression and genome stability depending on G4 localization^44^. In this regard, it is a remarkable coincidence that PRMT5 is upregulated in various cancer entities, while neoplasms are also known to have a high abundance of G4s, particularly in promoters driving oncogene expression^45^. Defining the regulatory impact of PRMT5 on G4-dependent processes might therefore have a broad relevance beyond the heart. Along this line, we identified Aurora kinase B (*AURKB*) as a gene targeted by G4-associated IR. Notably, therapeutic effects of PRMT5i were attributed to increased IR and subsequent loss of AURKB^46^, indicative of a G4-dependent splicing mechanism as in *ACADVL*.

### Loss of ACADVL is critical for PRMT5-dependent cardiomyopathy

Major transcriptional alterations were found in cardiac tissue upon deletion of *Prmt5*. One of the most altered pathways included FAO. Since altered energy metabolism has been described not only as a hallmark of advanced heart failure but also as a cause for cardiac dysfunction, we hypothesized that loss of ACADVL may drive progressive HFrEF. Indeed, our experiments showed that ACADVL is a critical downstream target of PRMT5 whose rescue is sufficient to prevent heart failure. This finding underlines the importance of intact lipid metabolism in adult cardiomyocytes^21^, which is also reflected by the clinical association between *ACADVL* loss-of-function mutations and severe infantile cardiomyopathy^10,47^. Given the broad range of enzymes and metabolic pathways affected by *Prmt5*-cKO, it is remarkable that loss of ACADVL alone is responsible for cardiac dysfunction, pointing to the central role of ACADVL in cardiomyocyte lipid metabolism.

Our findings raise the question if ACADVL deficiency-induced heart failure might be caused by an absolute deficit of energy-rich metabolites, or if the phenotype relates to secondary effects or non-canonical enzymatic properties. FAO represents the major source of energy in the adult heart^48^, using palmitic acid (C16:0) and oleic acid (C18:1) as the most abundant free fatty acids in the blood^49^. Because both fatty acids are substrates of ACADVL, their deficient degradation could cause an accumulation of toxic lipid species. Lipotoxicity in turn is known to negatively affect cardiac homeostasis^50^, while especially palmitate as a component of ceramides seems to be detrimental^51^. In this context, the repression of proinflammatory signaling by ACADVL, which appears key to its cardioprotective function, might be of particular relevance for the massive systemic increase of inflammatory chemokines in ACADVL deficiency (VLCADD) patients^52^. With the discovery of PRMT5- and G4-linked splicing control of ACADVL as a key metabolic enzyme in cardiomyocytes, targeted interventions to sustain this regulatory circuit may one day improve the treatment of heart failure.

## Acknowledgement

1. J. Krebs-Haupenthal, S. Harrack, M. Oestringer, S. Nazir, I. Broll, J. Poetzsch (all affiliated with the Institute of Experimental Cardiology, Medical Faculty, Heidelberg University, Germany) provided technical assistance. We thank M. Hagenmüller (Institute of Experimental Cardiology, Medical Faculty, Heidelberg University, Germany) for organizational support as well as G. Schmidt and S. Kaden (German Cancer Research Center, Heidelberg, Germany) for the help with light and electron microscopy. The authors further thank V. Benes and EMBL GeneCore (European Molecular Biology Laboratory (EMBL), 69117 Heidelberg, Germany) for support with sequencing and J. Hauke and K. Schwarz for metabolic measurements.

## Author contributions

JB and VM designed the study;

JB, CD and MD supervised the work, contributed to the data interpretation and provided financial resources.

VM, TM, JH, MTS, MD performed experiments;

VM, TBB, TM, EG, JH, MTS, FS, CUO, CS, JGO, AS, MD, CD, JB analyzed and interpreted data;

SS, JGO, GS, MWH, EEMF, AS, CD, JB provided research support and conceptual advice;

VM, TBB and JB wrote the manuscript with input from all authors.

## Supplementary Figures

Supplementary figure 1: PRMT5-controlled introns are spatially associated with G4s.

Supplementary figure 2: Loss of PRMT5 in cardiomyocytes induces heart failure.

Supplementary figure 3: *Acadvl*-AAV9 *in vivo*.

## Supplementary Tables

Supplementary table 1: List of candidates tested in the splicing screen

Supplementary table 2: Phenotypic characteristics of *Prmt5*-cKO in mice

Supplementary table 3: EdgeR and Majiq analysis results

## Data availability

Data sets were deposited at ArrayExpress and are accessible via E-MTAB-13594, E-MTAB-14401 and E-MTAB-14396.

Correspondence and requests for materials should be addressed to Johannes Backs.

## Methods

### Human myocardial samples

Myocardial tissue derived from patients suffering from end-stage dilated cardiomyopathy (DCM) was obtained within heart transplantation or implantation of assist devices. Control samples were obtained from non-failing donor hearts, which could not be transplanted for technical reasons. This set of samples used for immunoblotting was already published by Maack et al.^53^ and the corresponding study was approved by the Ethical Committee (Ärztekammer Saarland) and refers to the application 131/00.

### In vivo studies

All animal studies were conducted in accordance with the German Animal Welfare Act and approved by the Animal Experiment Review Board of the Regierungspräsidium Karlsruhe, Germany. The mouse studies refer to G124/16, G150/18 and G202/21.

### Generation of *Prmt5*-cKO mice

Mice harboring a conditional-ready allele of *Prmt5* were obtained from the European Mouse Mutant Archive (EMMA, B6Brd;B6N-Tyr^c-Brd^ A^tm1Brd^ Prmt5^tm2a(EUCOMM)Wtsi^Prmt5). To obtain a conditional KO allele the lacZ reporter was removed via a flippase recombinase which was subsequently outcrossed to obtain Prmt5^loxP/loxP^.

Cardiomyocyte-specific *Prmt5*-deleted mice were generated by mating Prmt5^loxP/loxP^ mice with transgenic mice expressing the inducible Cre recombinase alpha-myosin heavy chain promoter dependently (alpha-MHC-MerCreMer)^54^. All mouse lines were maintained in a C57BL/6N background. Genotyping was performed using following primer pairs:

alpha-MHC-MerCreMer:

For: 5’ ATA CCG GAG ATC ATG CAA GC 3’
Rev: 5’ AGG TGG ACC TGA TCA TGG AG 3’
For: 5’ CTA GGC CAC AGA ATT GAA AGA TCT 3’
Rev: 5’ GTA GGT GGA AAT TCT AGC ATC ATC C 3’

Prmt5-loxP:

For_1: 5’ CTG CAC ACA CAT GGC ACA TAT ACA G 3’
For_2: 5’ TGG AAC TGC AGG CAT ATG CCA C 3’
Rev: 5’ GGC AAG AAC ATA AAG TGA CCC TCC 3’

The genetic deletion via Cre recombinase activation was induced at the age of 8 weeks via oral gavage of Tamoxifen (Sigma) over 2 cycles each 5 subsequent days with a resting time of 2 days in between. Tamoxifen was dissolved in sunflower oil and 6% Ethanol to obtain a 10 mg/ml stock solution. The final dosage administered to the mice once a day was 75 mg per kg body weight.

After Tamoxifen application, an interval of 3 weeks was kept for recovery and prevention of Tamoxifen-dependent side effects until Isoprenaline treatment was started. Cre-positive (Cre^+/-^) *Prmt5* wildtype animals in the same genetic background served as controls. Since no sex-dependent differences of the phenotype were observed, male and female littermates were used in parallel.

## Animal procedures

### Transthoracic echocardiography

To monitor cardiac performance transthoracic echocardiography was performed. Mice were shaved and stayed conscious during the examination. Measurements were assessed using a Vevo® 2100 system with a 55 MHz transducer (Visualsonics). Ejection fraction (EF) was averaged over at least 5 contraction cycles. The experimenter was blinded for genotype and treatment.

### Isoprenaline treatment

Isoprenaline (Sigma) was dissolved in 0,9% NaCl supplemented with 0,1% Ascorbic acid (Roth). It was administered intraperitonally once a day at a final dosage of 30 mg per kg body weight for 3 or 10 subsequent days, respectively. 24h after the last application mice were euthanized by cervical dislocation and organs were harvested.

## Generation and application of AAV9 vectors

Cardiotrophic AAV9 vectors were generated as described elsewhere^55^. In brief, protein coding cDNA sequence of murine *Acadvl* transcript ENSMUST00000102574.10 was cloned into a pSSV9-TnnT vector enabling cardiomyocyte-specific expression controlled by the troponin T (Tnnt) promoter ^56^. An AAV9 construct encoding renilla luciferase (pSSV9-TnnT-Luc) was used as control. Packaging into AAV9 capsids was achieved by co-transfection of genomic plasmids and pDP9rs in HEK293T cells. Cell lysates and supernatant were subsequently used for virus purification and final quantification was performed via qPCR.

The AAV9 vectors were injected into the tail vein at indicated dosages. Prior to the main study a dose-finding trial was conducted to evaluate appropriate dosing for compensative *Acadvl* expression. A dosage of 3×10e+11 viral genomes of *Acadvl*-AAV9 and luciferase-AAV9 was used for final studies, respectively.

## Lipidomics

Lipids were extracted from murine left ventricular tissue. Heart tissue was homogenized using a Retsch mill and ceramic beads in 0.5 mL ice-cold MeOH/H2O (80/20 v/v)/10mg tissue. The suspension was transferred to a glass tube and another 0.5 ml ice-cold MeOH/H2O (80/20 v/v) were added. Internal standard mix (SPLASH LIPIDOMIX (Avanti: 330707-1EA)) was used for normalization 5uL/10mg tissue. After addition of 120 µl 0.2 M HCl, 360 µl chloroform, 400 µl chloroform and 400 µl of water with vigorous mixing between the pipetting steps, samples were centrifuged at 3,000g for 10 min. 700 µl of the lower lipid phase was collected and taken to dryness under a stream of nitrogen gas at 40°C. Lipids were then dissolved in 200 µl isopropanol and subjected to LC-MS analysis. Lipids were separated on a C8 column (Accucore C8 column, 2.6 µm particle size, 50 x 2.1 mm, Thermo Fisher Scientific) mounted on an Ulitmate 3000 HPLC (Thermo Fisher Scientific) and heated to 40°C. The mobile phase buffer A consisted of 0.1% formic acid in CH3CN/H2O (10/90, v/v) and buffer B consisted of 0.1% formic acid in CH3CN/IPOH/H2O (45/45/10, v/v/v). After injection of 3 µl lipid sample, 20% solvent B were maintained for 2 minutes, followed by a linear increase to 99.5% B within 5 minutes, which was maintained for 27 minutes. After returning to 20% B within 1 minutes, the column was re-equilibrated at 20% B for 5 minutes, resulting in a total run time of 40 minutes. The flow rate was maintained at 350 µl/min and the eluent was directed to the ESI source of the QE Plus from 2 to 35 minutes. MS analysis was performed on a Q Exactive Plus mass spectrometer (Thermo Fisher Scientific) applying the following settings: Scan settings: Scan range – 200-1600 m/z in full MS mode with switching polarities (neg/pos) and data-dependent fragmentation; Resolution – 70,000, AGC target – 1E6; Max. injection time – 50 ms. HESI source parameters: Sheat gas – 30; Aux gas – 10; Sweep gas – 3; Spray voltage – 2.5 kV; Capillary temperature – 320°C; S-lens RF level – 55.0; Aux gas heater temperature – 55°C; Fragmentation settings: Resolution – 17,500; AGC target – 1E5; Max. injection time –50 ms Peaks corresponding to the calculated lipid masses (±5 ppm) were integrated using El-Maven (https://resources.elucidata.io/elmaven).

## Acyl-Carnitine profiling

Murine left-ventricular tissue was homogenized in sterilized water. Samples were centrifuged and resulting supernatants were extracted like serum and the acylcarnitines were analyzed by electrospray ionization tandem mass spectrometry (ESI-MS/MS) according to a method as previously described^57^, using a Waters Xevo TQDQuattro Ultima triple quadrupole mass spectrometer (Waters GmbH, Eschborn, Germany) equipped with an electrospray ion source and a Micromass MassLynx data software (Micromass, Manchester, UK). Results were normalized to total protein content.

## Cardiomyocyte-specific nuclei isolation

Isolation of cardiomyocyte-specific nuclei was performed as described elsewhere^58^ including the following adaptations. Cardiomyocyte nuclei were stained with anti-PCM1 (Sigma HPA023374) primary and Alexa405-labled secondary antibody (Thermo Fisher A48258) and sorted on a Sony SH800 system. Unspecific nuclear staining was achieved via Draq7 (Thermo Fisher D15105). 10.000 PCM1-positive nuclei per biological replicate were subjected to RNA extraction based on the RNeasy Micro Kit (Qiagen) and on-column DNAse treatment.

## In vitro studies

Cell culture: HEK293 cells (ATCC) were maintained in DMEM (PanBiotech) supplemented with 10% FCS (Thermo Fisher) and 1% penicillin-streptomycin (Thermo Fisher) and incubated at standard conditions (humidified, 5% CO2/95% room air, 37°C).

EPZ015666 (MedChemExpress) was dissolved in DMSO and added into the culture medium at 0.1 to 10µM for 96h (protein analysis, Fig. 2B). For subcellular RNA analyses and longread sequencing cells were treated with 1µM EPZ015666 for 48h.

Pyridostatin (Medchemexpress) was dissolved in water or PBS and added into the culture medium at dosages between 1µM and 100µM for 24h (RNA analysis) or 10µM for 48h (protein analysis).

Taq polymerase stalling was performed according to manufacturer’s recommendations at dosages between 10nM and 100nM. 7-deaza-dGTP (NEB) was used at equivalent dosages as dGTP in the reaction mix.

Primer:

mmAcadvl_E17 for: GGCTCTGGATCAATTTGCCAC
mmAcadvl_E18 rev: GAACCACCACCATGGCATAGA
mmAcadvl_E14 for: AGCGTGTGCTCCGAGATATT
mmAcadvl_E15 rev: CCTTTCCTTTGTCCATACAGCC

To determine splicing behavior of a minigene construct, protein coding cDNA sequence of murine *Acadvl* transcript ENSMUST00000102574.10 including intron 17/18 (chr11:70,010,771-70,010,848 (GRCm38)) or intron 8/9 (chr11:70,013,084-70,013,171) or a G4 mutant (point mutations in intron 17/18 correspond to Fig. 1E) were cloned into a pcDNA vector. Followed by a pre-treatment with EPZ015666 and DMSO, respectively, for 24h, the constructs was transfected into HEK293 cells using GeneJammer (Agilent, 204130) according to manufacturer’s protocol and nuclear RNA was isolated after 24h. The empty pcDNA vector served as control.

Knockdown of candidate proteins (Suppl. Table 1) was performed using the RNAi Consortium (TRC1 human shRNA library), specifically:

TRCN0000006630 and TRCN0000006631 targeting *SRSF9*
TRCN0000074474 and TRCN0000074475 targeting *WBP11*
pLKO.1-Scrambled (Addgene #136035)

Constructs were transfected into HEK293 cells using GeneJammer followed by Puromycin selection (Sigma, P8833) at 2µg/ml and nuclear RNA isolation or protein extraction, respectively.

Myc-tagged constructs of human SRSF9 and WBP11 were obtained by cloning protein coding cDNA sequences (SRSF9: ENST00000229390.8; WBP11: ENST00000261167.7) along with a C-terminal myc into a pcDNA vector. Transfection into HEK293 cells was performed using PEI MAX (Polysciences).

Seahorse experiments were performed and analyzed according to manufacturer’s recommendations in HEK293 cells upon pre-treatment with 1µM EPZ015666 for 48h. Cells were starved for 6h and exposed to palmitate-BSA conjugate or BSA control (Cayman Chemical) immediately before starting the assay.

## G4 detection

Thioflavin T staining of DNA G4 oligonucleotides (Eurofins Genomics) was performed as published earlier^59^:

Transcribed strand:

CTGTCAGGTAGATGAGGGTGGGGCTGTTAGTCGGAGGGAGGGAATCAGGGAGGGAAGGGGGTTGCAGAGCTCACTCACTGACAATCCCTTTCTTGTGTTTCAC

Non-transcribed strand:

GTGAGTGAGCTCTGCAACCCCCTTCCCTCCCTGATTCCCTCCCTCCGACTAACAGCCCCACCCTCATCTACCTGACAGACGAGCAGTTCCTGCTGCA

Transcribed strand G4 mutant:

GTGAGTGAATCAGTGAGTGAAGTGTGTTGCAGAGCTCACTCACTGACAATCCCTTTCTTGTGTTTCAC

Non-transcribed strand G4 mutant:

GTGAGTGAGCTCTGCAACACACTTCACTCACTGATTCACTCACTCCGACTAACAGCCCCACCCTCATCT

G4-specific nanobody SG4 and the corresponding R105A mutant were produced and purified as published^19^. Lable-free Electrophoretic mobility shift assay (EMSA) was performed in binding buffer (20mM Tris-HCl pH 8.0, 50mM NaCl, 80mM KCl, 8% glycerol) using 1µg protein^60^ and the gel was stained with SYBR Gold (Thermo Fisher). DNA oligonucleotide sequences correspond to the ThT assay.

Streptavidin-mediated pulldown of biotinylated G4 oligos was performed using lysates of HEK293 cells in a buffer containing 20mM Hepes-KOH pH 7.9, 100mM KCl, 0.5mM EDTA, 10% Glycerin, 1mM DTT supplemented with protease inhibitor. Binding reaction was performed for 1,5h at 5°C in 20mM Hepes pH 7.4, 50mM KCl, 0.01% NP40, 0.5mM EDTA using the entire wildtype or G4-mutated DNA sequence of intron 17/18, respectively (Eurofins Genomics), pre-coupled to Dynabeads MyOne Streptavidin C1 beads (Invitrogen). Bound proteins were analyzed via SDS-PAGE and Western Blotting, or Coomassie staining (Serva).

## Immunoprecipitation (IP)

HEK293 nuclei were lysed in IP buffer (10% glycerol, 1% Triton X-100, 1,5 mM MgCl2, 50 mM HEPES-NaOH, pH 7.5, 150 mM NaCl, 1 mM EDTA), supplemented with protease inhibitor. Pre-incubation of lysate and respective antibodies was performed overnight and antigen-antibody complexes were precipitated using Protein G Dynabeads. Proteins were eluted in Laemmli sample buffer upon heating to 95°C.

## Immunoblotting

Human and murine heart tissues were homogenized in Kranias buffer (30mM Tris-HCl pH 8,8, 5mM EDTA, 30mM NaF, 3% SDS, 10% Glycerol) containing protease and phosphatase inhibitor cocktail (Sigma). Protein lysates were separated by SDS-PAGE and blotted onto nitrocellulose membranes (Amersham) followed by blocking with Roti-Block (Roth) and overnight incubation with primary antibodies. Subsequently, membranes were washed and probed with secondary antibodies. Chemiluminescence was measured using a Fusion FX system (Vilber).

Primary antibodies:

Anti-PRMT5 antibody (Merck/Millipore, 07-405)
Anti-Dimethyl-Arginine antibody, symmetric SYM10 (Merck/Millipore, 07-412)
Anti-ACADVL antibody (Santa Cruz Biotechnology, sc-376239)
Anti-SRSF9 antibody (Thermo Fisher Scientific, TA808492)
Anti-WBP11 antibody (Thermo Fisher Scientific, PA5-31241)
Anti-ILF3 antibody (Abcam, ab131004)
Anti-myc antibody (Merck/Millipore, 05-724)
Anti-flag antibody (Sigma-Aldrich, A8592)
Anti-GAPDH antibody (Merck/Millipore, MAB374)
Anti-beta tubulin antibody (Sigma-Aldrich, T4026)
Anti-histone 3 antibody (Sigma-Aldrich, H9289)

Secondary antibodies:

Goat Anti-Mouse (SouthernBiotech, 1031-05)
Goat Anti-Rabbit (SouthernBiotech, 4050-05)

## RNA extraction and quantification

Tissue and cells were lysed in QIAzol (Qiagen) or based on the RNeasy Mini Kit (Qiagen) according to manufacturer’s protocol.

For subcellular RNA isolation cells were lysed in buffer containing 0,1% Tween20 and 0,1% NP40 supplemented with β-Mercaptoethanol (Sigma) or RNasin Plus (Promega). Nuclei were subsequently pelleted by centrifugation for 5min at 600g. The supernatant was kept as cytosolic fraction and both fractions were further subjected to RNA extraction.

1µg of total RNA was reverse transcribed using the PrimaDIRECT 2x PCR Master Mix Kit (Steinbrenner). Quantitative PCR was performed on a LightCycler480 instrument (Roche) using SYBR Green Master Mix (Thermo Fisher).

Primer sequences used for quantitative PCR:

hsAcadvl_spliced:

For: 5’ TGG TGG AGG CCA AGC TGA TAA 3’
Rev: 5’ GAG AGA ACC ACC ACC ATG GCA TA 3’

hsAcadvl_intron:

For: 5’ TGG TGG AGG CCA AGC TGA TAA 3’
Rev: 5’ AAA GGG AGG GGC GGA ATG G 3’

hsGapdh:

For: 5’ ACC CAC TCC TCC ACC TTT GAC 3’
Rev: 5’ ACC CTG TTG CTG TAG CCA AAT T 3’

mmAcadvl_spliced:

For: 5’ GGC TCT GGA TCA ATT TGC CAC 3’
Rev: 5’ ACT GCT CGT TGA CAA TCC CTT 3’

mmAcadvl_intron:

For: 5’ GGC TCT GGA TCA ATT TGC CAC 3’
Rev: 5’ TAG ATG AGG GTG GGG CTG TT 3’

mmAcadvl total:

For: 5’ TAC CCA GTA TGC TCG CTT GG 3’
Rev: 5’ CTG GGC CTT TGT GCC ATA GA 3’

mmCxcl10:

For: 5’ CCG TCA TTT TCT GCC TCA TCC 3’
Rev: 5’ CTG CTC ATC ATT CTT TTT CAT CGT G 3’

mmCcr5:

For: 5’ GGA GGT GAG ACA TCC GTT CC 3’
Rev: 5’ GAG CTG AGC CGC AAT TTG TT 3’

mmGapdh:

For: 5’ GGT GGA CCT CAT GGC CTA CA 3’
Rev: 5’ CTC TCT TGC TCA GTG TCC TTG CT 3’

## G4 motif conservation in mammals

Depicted sequences correspond to:
Mus musculus: intron 17 (ENSMUST00000102574.10)
Homo sapiens: intron 17 (ENST00000356839.10)
Sus scrofa: intron 16 (ENSSSCT00035082964.1)
Oryctolagus cuniculus: intron 17 (ENSOCUT00000006970.4)
Gorilla gorilla: intron 19 (ENSGGOT00000006789.3)
Balaenoptera musculus: intron 15 (ENSBMST00010033027.1)
Canis lupus familiaris: intron 19 (ENSCAFT00805007598.1)
Loxodonta africana: intron 17 (ENSLAFT00000018642.4)

All guanine repeats were highlighted including predicted G4 motifs based on pqsfinder^61^.

## RNA sequencing and bioinformatical analysis

RNA sequencing was performed on murine left ventricular tissue after a 3-days-treatment of ISO or control saline solution. Library preparation and sequencing as 100 bp paired-end (BGISEQ-500) was performed by BGI, Hong Kong.

Nuclear RNA sequencing was performed according to the SmartSeq2 protocol^62,63^. 200pg cDNA were used for tagmentation and libraries were sequenced with NextSeq2000 100 bp paired-end.

Both left-ventricular tissue and CM nuclear RNA sequencing were processed with the same computational workflow. First, low-quality bases were removed with Flexbar (v3.5.0)^64^. Next, short reads from the rRNA loci were subtracted by mapping against the Mus musculus ribosomal RNA precursor using Bowtie2 (v2.4.5)^65^ and then discarded. The remaining reads were aligned to the Mus musculus genome (Ensembl transcriptome Annotation GRCm38 v102) with STAR (v2.6.0c)^66^. Gene counts were quantified with StringTie (v2.2.1)^67^. Differential gene expression analysis was conducted with EdgeR with the quasi-likelihood negative binomial generalized log-linear model. The LV tissue and nucCM were analyzed with the ∼ Treatment + Genotype, ∼ Treatment + Genotype + Sex + Batch design, respectively. Genes with adjusted p-value < 0.05 and log_2_ fold change > 1 were called significant. Gene set enrichment analysis was obtained with FGSEA (v1.22.0)^68^ with WikiPathways ontology (release 20240210) and terms with adjusted p-value < 0.01 were considered significant. Differential splicing analysis was conducted with Majiq (v2.2-e25c4ac)^69^ via the Baltica workflow (v1.2.4)^70^ using the Majiq deltaPSI threshold of 0.20. Splicing events with absolute deltaPSI > 0.2 and probability no-change < 0.05 were called significant.

To analyze the association between G4 structures^16^ and intron retention, G4 coordinates originally in the hg19 assembly were first converted to hg38 using UCSC’s liftOver tool^71^, ensuring compatibility with IRFinder-derived intron annotations. Bedtools-v2.29.2 intersect^72^ was then employed to identify overlaps between G4 regions (±0, 50, 100, 200, 350 and 500 bp windows) and introns, stratified by strand. Stratification by transcriptional direction was in this case performed by using GENCODE Human Release 48^73^. Nested introns were removed to avoid counting overlaps multiple times and we ensured that each intron contributed only once to the counts. Enrichment percentages were calculated per condition and distance window, and significance between retained and non-regulated overlaps was assessed using proportion tests.

Genome-wide non-canonical structure analysis was performed based on the Non-B database v2.0^74^ mapped to hg38 assembly. Significantly retained introns (padj <= 0.05) were used for enrichment analyses along with different spatial extensions (200bp, 500bp) beyond intron junctions. A Fischer’s test was used to report for the odds ratio (OR).

## Statistical analysis

Data sets are displayed as mean ± SEM. GraphPad Prism 9 software was used for statistics and visualization. Statistical testing was performed by either One-Way ANOVA and Tukey’s multiple comparisons test with multiplicity adjusted P values or unpaired t-test. If not otherwise indicated, p<0.05 was considered as significant.

---

**Supplementary Figure 1:**
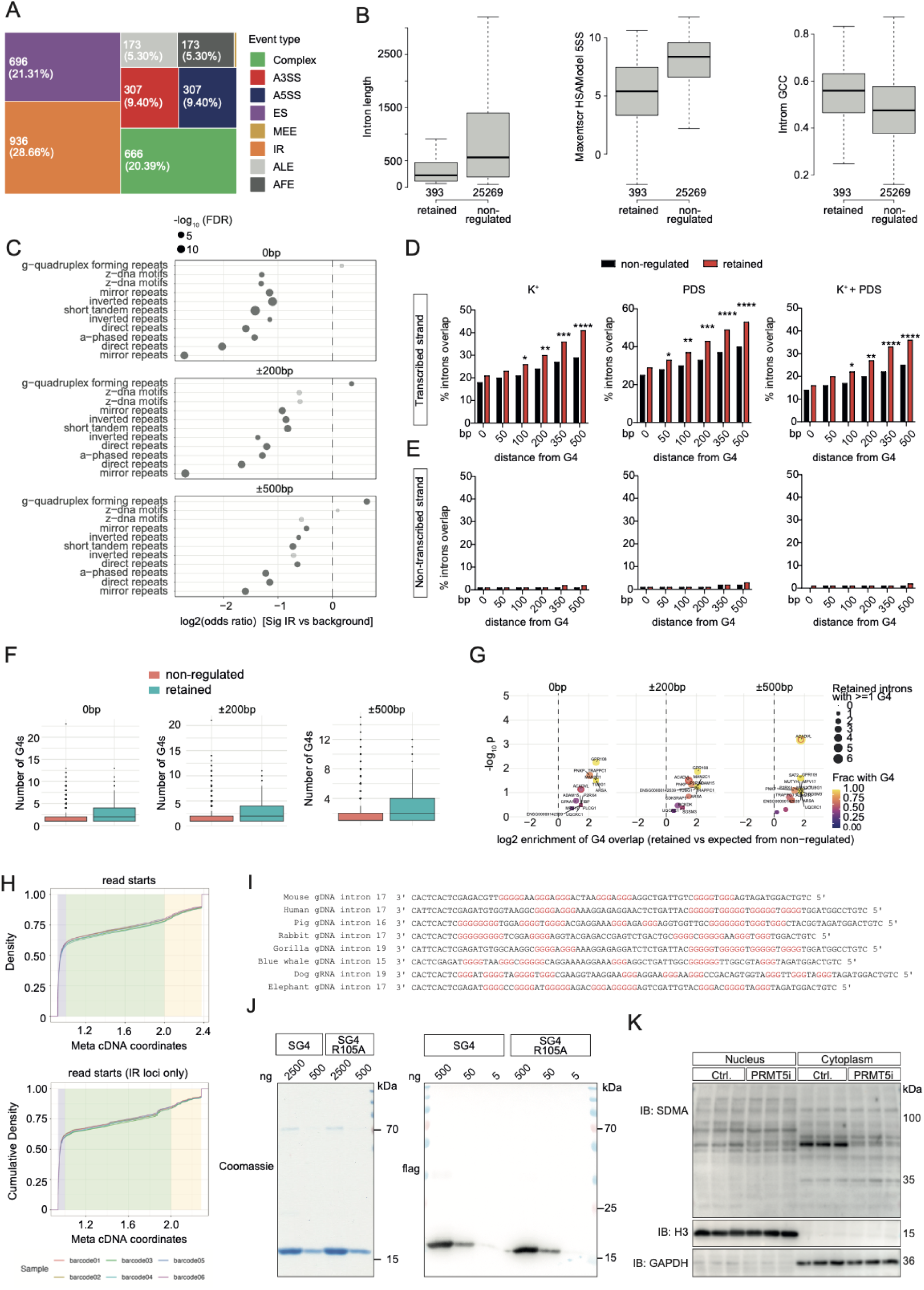
PRMT5-controlled introns are spatially associated with G4s. (**A**) Size-proportional representation of absolute numbers of alternative splicing events in HEK293 nuclei. We distinguish intron retention (IR), exon skipping (ES), alternative 3‘ splice site (A3SS), alternative 5‘ splice site (A5SS), mutual exclusive exons (MEE), alternative last exon (ALE), alternative first exon (AFE) as well as complex events. (**B**) General characteristics of significantly retained compared to non-regulated introns including intron length (left), 5’ splice site strength (middle) and intronic GC content (right). (**C**) Non-canonical structure prediction reveals strong enrichment of G4s and significantly retained introns upon PRMT5i on a genome-wide basis across window sizes between 0bp and 500bp beyond intron junctions. Bold dots represent FDR<0.05. (**D, E**) Integrated analysis of G4-seq and ONT-seq in HEK293 nuclei upon PRMT5i across different experimental conditions to stabilize G4s (potassium ions, Pyridostatin (PDS) or a combination of both). The correlation of retained and non-regulated introns is shown for different windows around the G4 (x axis). Transcribed and non-transcribed DNA strand are separately analyzed. Statistical analysis: K^+^: * p=0.0458, ** p=0.00621, *** p=0.00012, **** p<0.0001, ** p=0.00203; PDS: * p=0.0391, ** p=0.00486, *** p=0.000102, **** p<0.0001; K^+^ + PDS: * p=0.0355, ** p=0.00116, **** p<0.0001 (Proportion test). (**F**) Genome-wide correlation analysis between genomic G4s on the transcribed strand and retained introns. Introns were first grouped based on PRMT5-dependent splicing outcome. Local abundance of known G4s were counted with and without windows as indicated. Statistical analysis: **** p=4.654*e^-14^ (0bp), **** p=9.048*e^-16^ (200bp), **** p=2.922*e^-15^ (500bp) (Wilcoxon rank sum test). (**G**) Extended G4 enrichment analyses on gene level highlights *ACADVL* as prominent target of G4-associated IR, particularly when applying a window of 500bp beyond intron junctions indicating significant correlation of DNA G4s and IR. (**H**) Meta cDNA plot of Nanopore long-read sequencing reveals no significant difference in read density between Ctrl. and PRMT5i. (**I**) Comparative depiction of intronic sequences across different mammals. Guanine repeats are highlighted. Each gDNA sequence is predicted to form a G4. (**J**) Coomassie staining and anti-flag immunoblot analysis of the recombinant nanobody used for EMSA. (**K**) Immunoblot analysis of subcellular fractions of HEK293 cells treated with PRMT5i. GAPDH and histone 3 (H3) served as fraction-specific controls.

**Supplementary Figure 2:**
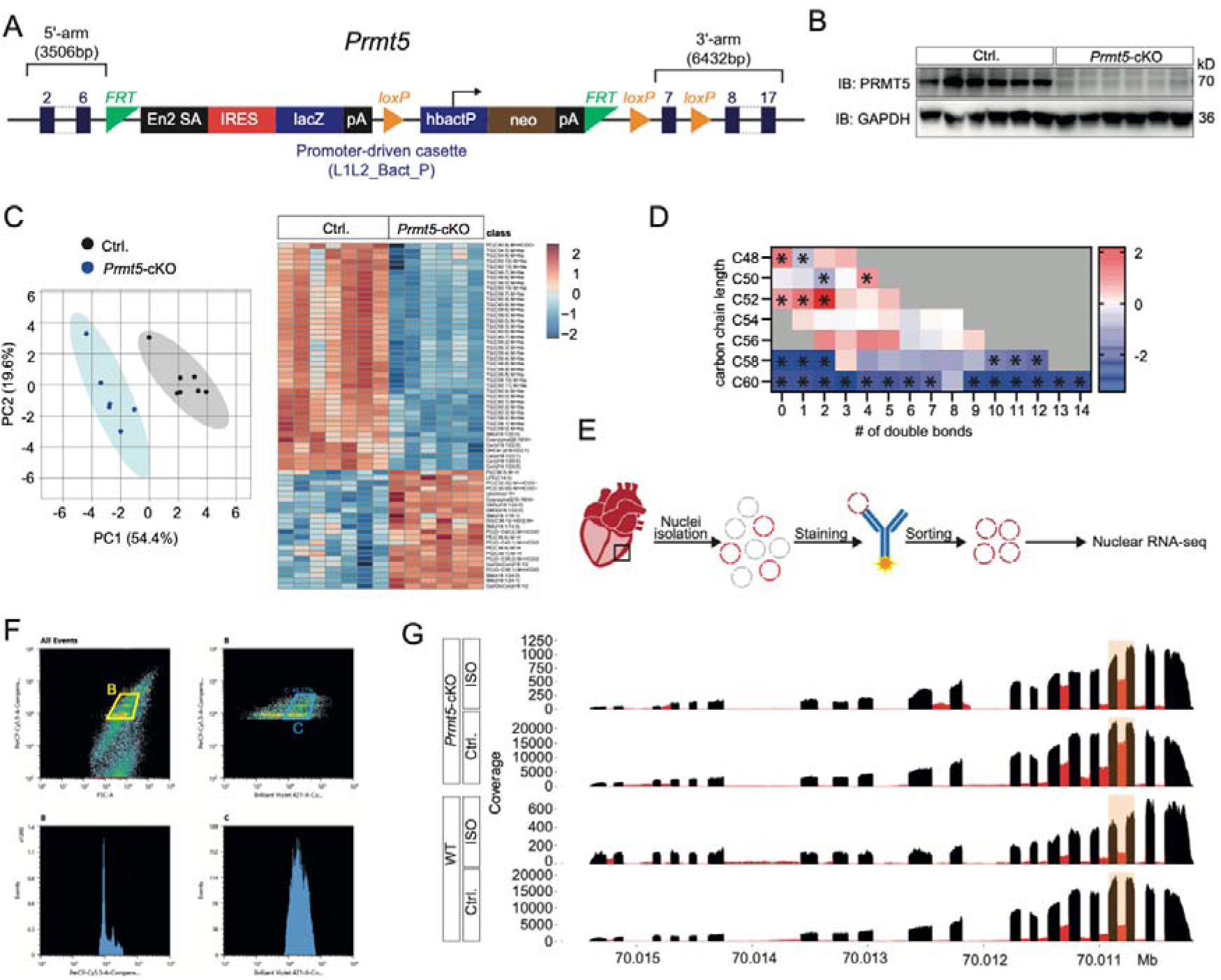
Loss of PRMT5 in cardiomyocytes induces heart failure. (**A**) Scheme on genetic knockout strategy used for generating *Prmt5*-cKO mice. (**B**) Immunoblot analysis of PRMT5 in cardiac tissue of *Prmt5*-cKO and control mice (Ctrl.). GAPDH served as loading control. (**C**) Left: Principal component analysis of lipidomic profiling of cardiac tissue derived from aged *Prmt5*-cKO and Ctrl. mice. Right: Analysis of cardiac lipidome in aged mice. Normalized abundance is displayed as Z-score. (**D**) Subanalysis of triglyceride (TG) fraction according to carbon chain length and number of double bonds. Abundance is displayed as log2-FC. Significantly regulated entities with p adj. < 0.05 are marked by asterisks. (**E**) Schematic depiction of the workflow to isolate CM-specific nuclei from heart tissue including nuclei isolation, fluorescence staining and sorting of PCM1-positive nuclei which were subsequently subjected to RNAseq. (**F**) Representative fluorescence-activated nuclei sorting (FANS) using Draq7 as nuclear staining (B) and PCM1 for CM specificity (C). (**G**) Integrated mean coverage plot of the *Acadvl* locus for *Prmt5*-cKO and Ctrl. with n≥3. Conditions are identical to Fig. 4L.

**Supplementary Figure 3:**
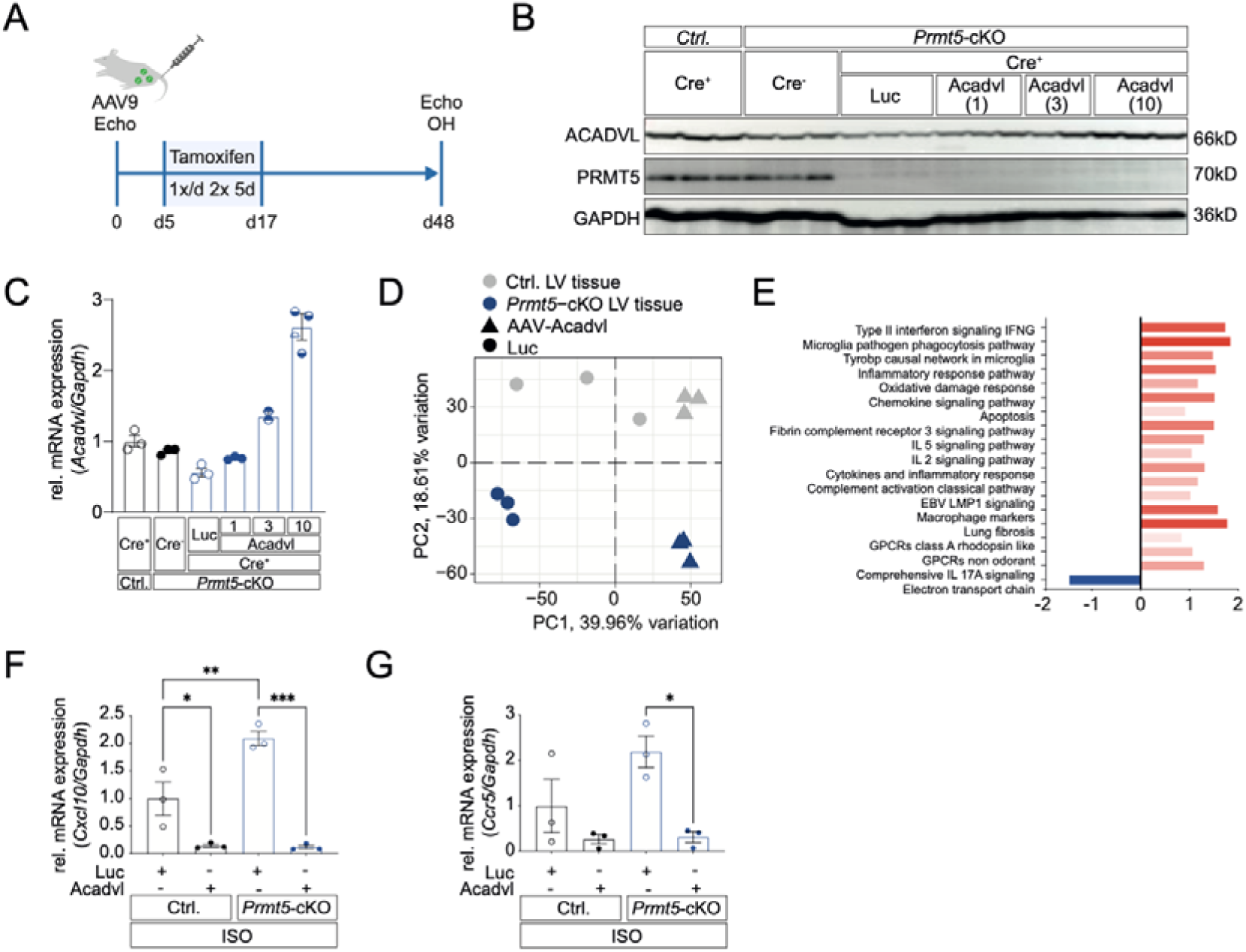
*Acadvl*-AAV9 in *vivo*. (**A**) Experimental course of the dose finding study including AAV9 injection, KO induction via Tamoxifen and organ harvest after 48 days (d). (**B**) Immunoblot analysis of ACADVL and PRMT5 in LV tissue of *Prmt5*-cKO and control mice (Ctrl.) 48 days after AAV injection. Dosages covered 1×10e+11 (1), 3×10e+11 (3) and 1×10e+12 (10) viral genomes. As control for endogenous ACADVL expression Cre^+^ Ctrl. as well as Cre^-^ *Prmt5*-cKO animals were used. GAPDH served as loading control. (**C**) Quantitative PCR on *Acadvl* in LV tissue of *Prmt5*-cKO and control mice (Ctrl.). Individual samples and conditions are identical to (**B**). Values were normalized to *Gapdh* and are presented as Mean±SEM. (**D**) Principal component analysis on RNAseq of left-ventricular (LV) tissue of PRMT5-deficient and control (Ctrl.) mice after Acadvl-AAV and luciferase (Luc)-AAV treatment and 10 days ISO stimulation. (**E**) Pathway analysis of RNA sequencing of cardiac tissue of *Prmt5*-cKO compared to Ctrl. samples (genotype coefficient) after 10 days of ISO stimulation. Pathways correspond to Fig. 5E. (**F, G**) Quantitative PCR on *Cxcl10* (**F**), and *Ccr5* (**G**) in left-ventricular tissue of AAV-treated mice. Values were normalized to *Gapdh*. Statistical analysis: values are presented as Mean±SEM (**F**) * p=0.0259, ** p=0.0071, *** p=0.0001 (Ordinary one-way ANOVA and Tukey‘s multiple comparisons test) (**G**) * p=0.0217 (Ordinary one-way ANOVA and Tukey‘s multiple comparisons test).

